# Trends Assessment of the Genetic Mutation Induced Protein-Protein Interaction Variation via Protein Large Language Driven Method

**DOI:** 10.1101/2025.05.23.655070

**Authors:** Zhejun Kuang, Yunkai Li, Zhe Liu, Jian Zhao, Lijuan Shi, Akira-Kawai, Han Wang

## Abstract

Protein-protein interaction (PPI) variations are widely observed in many principal biological processes related to various diseases, while genetic mutation is one of the most common factors resulting in those variations. Unraveling these underlying biomolecular interactions is a key challenge that hinders a profound understanding of disease mechanisms. Although current models can accurately predict quantitative binding affinity, their application in pathogenic research is limited by difficulties in defining trend types, particularly extreme “disrupting” cases, along with poor generalizability beyond single-point mutations (SNP) and insufficient biological interpretability. Here, we present TAPPI, an interpretable large language model-driven deep learning framework, to perform Trends Assessment of Protein-Protein Interaction. It categorizes PPI variations’ trends into four types: disrupting, decreasing, no effect, and increasing, enabling fine-grained functional assessments and bridging genetic mutation and disease mechanisms. The framework generalizes effectively across both single-point and multi-point mutations and captures complex trend outcomes relevant to diverse disease landscapes. In benchmark evaluations, TAPPI achieved the state-of-the-art prediction performance and demonstrated biological relevance through interpretable results. In external datasets from Autism Spectrum Disorder, Lennox-Gastaut syndrome, and Epileptic Encephalopathies, given the premise that TAPPI’s predicted results align with pathogenic patterns, these results reveal how genetic mutations indirectly perturb disease-associated pathways and supporting our perspective of “crucial disrupting” and “bridging mutations-disease”. In summary, accurate, interpretable, and pathogenically relevant prediction of PPI variations trend predicted by TAPPI provides a novel approach for mechanistic discovery.

## 1. Introduction

Protein-Protein interactions (PPI) variations are closely related to cellular function and stability [1, 2]. Alterations in amino acid sequences caused by single nucleotide polymorphisms (SNPs) and de novo mutations is one of the most common factors resulting in those variations, subsequently affecting the stability of keg pathway [3, 4] and increasing the risk of disease [5–8]. Assessment these PPI variations trend is essential for understanding the complex relationships within cellular processes [9, 10], revealing how they perturb normal biological pathways and lead to disease [11, 12], bridging mutation and disease.

Since protein binding affinity is the direct way to elucidate PPI variants, many efforts have focused on the corresponding prediction and have made some progress, such as MuToN [16] achieve state-of-the-art performance than mCSM-PPI2 [17], GeoPPI [18] and so on. Three inputs are needed for MuToN when predicting the PPI variation trend represented by the change in Gibbs free energy (ΔΔG): the spatial structure and sequence of the protein before mutation, the spatial structure and sequence after mutation, and the spatial structure and sequence of the binding protein. However, challenges in categorizing trend types in intricate cellular contexts, especially in extreme ‘Disrupting’ cases, along with reliance on structural information, restrict research on pathogenesis.

Liu et al. developed MIPPI [19] to assess PPI variation through a four-category classification trend (increasing, decreasing, disrupting, and no effect), differing from ΔΔG, providing a more straightforward and simplified approach to clinical assistance. Meanwhile, MIPPI solely uses protein sequences and evolutionary conservation information to perform categorical PPI variation trend prediction based on the sequences near mutation sites and the binding protein sequences for only single-point mutations. Despite the achievements mentioned before, it is still a challenge to achieve substantial prediction accuracy for binding affinity.

In this study, we propose TAPPI (Trends Assessment of Protein-Protein Interaction), a protein Large Language Model (LLM) driven deep learning framework, utilizes LLMs and improved backbones to achieve accurate predictions of the straightforward and simplified four classifications of PPI variation trends (increasing, decreasing, disrupting, and no effect) caused by both single-point and multi-point mutations without relying on structural information, effectively bridging gene mutations and disease. In benchmark evaluations, we demonstrated TAPPI’s predictive capabilities for both single-point mutations and short-distance multi-point mutations caused trends, achieving state-of-the-art performance. Additionally, we showcased TAPPI’s ability in predicting PPI variation trend caused by reverse mutations, its distribution in encoding space, and interpretability, confirming that the predictions adhere to biological norms.

In external datasets from Autism Spectrum Disorder, Lennox–Gastaut syndrome, and Epileptic Encephalopathies from the PsyMuKb database [5], we highlight that “In the PPI network, the PPI variation trends involving partner proteins closer to disease-related proteins show a higher proportion of ‘Disrupting’ and a lower proportion of ‘No_Effect’ in patients”. This emphasizes the accuracy of TAPPI’s predictions and suggests that mutations can indirectly contribute to disease through PPI variation. We then present the enrichment of partner proteins associated with PPI variation trends categorized as ‘With Effect’ (increasing, decreasing, and disrupting) in disease-related pathways for these patients and use cases companion with literatures to illustrate how mutations directly and indirectly influence disease-related pathways through PPI variation, demonstrates the potential of TAPPI to bridge mutations and disease, as well as its ability to assist in precise diagnosis at the individual level. Moreover, we highlighted TAPPI’s potential in identifying key mutations when examining multiple long-distance mutations.

In summary, TAPPI combines accuracy, interpretability, and biological relevance, making it a powerful tool for studying disease mechanisms at both population and individual levels. It uniquely uncovers how mutations indirectly influence pathogenic pathways through PPI variations, bridging mutations and disease phenotypes.

## 2. Result

### 2.1 Overview of TAPPI

TAPPI was designed to predict the impact of variation on protein-protein interaction (PPI). To achieve this, we collected 30,173 entries from the IMEx Consortium [20] after adding reverse mutations to enhance real-world relevance. All data were human-curated from published references, we categorized them into four main classes based on original labels: disrupting, decreasing, no effect, increasing, allowing TAPPI to give a straightforward and biological relevance prediction of trend on PPI variation.

We retained single-point mutations and multi-point mutations with a maximum distance of 20 amino acids, which corresponds to the size of the receptive field, rather than relying solely on single-point mutations. This approach allows TAPPI to learn information from sequences near the mutation without being overly dependent on the center of the receptive field and can handle multi-point mutations.

TAPPI’s architecture comprises two core components: (1) We use both local features of the 20 amino acids surrounding the mutation and global features of the whole protein extracted by protein-Large-Language-model (ESM-2) [21] to represent mutations. (Fig. 1b) Features of each amino acid and the result feature vectors [22] are used to represent the partner protein (Fig. 1c). (2) We employ multiple stacked cross-attention layers to enable the change of interaction between two proteins (Fig. 1d).

**Figure 1.**
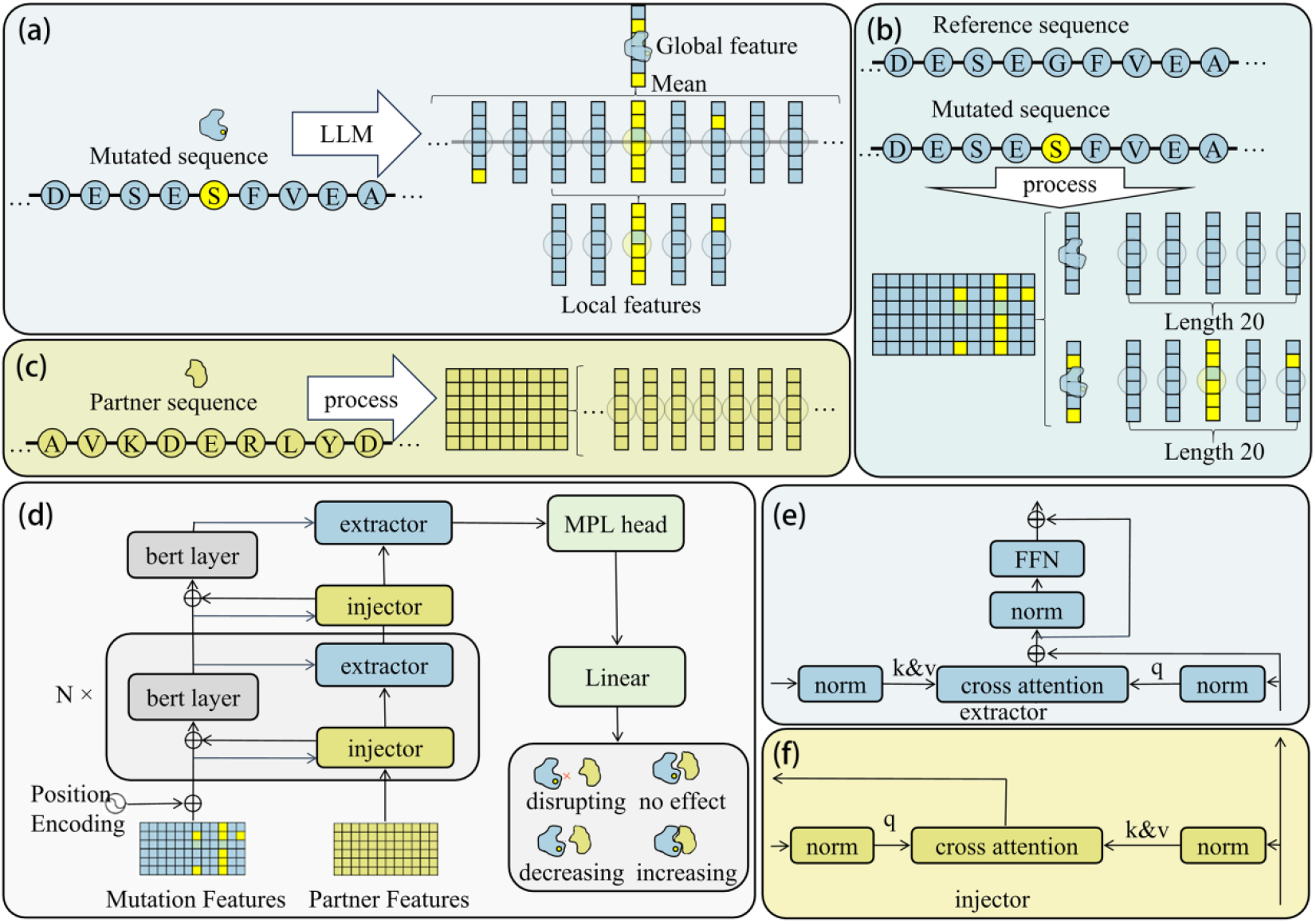
This figure illustrates the research workflow of TAPPI. (a) For the amino acid sequences before and after the protein mutation, the features of each amino acid are extracted using protein large language model (ESM-2). The features of amino acids near the mutation site are considered local features, while the average of all amino acid features represents the global features. (b) The local and global features of the amino acids before the mutation are concatenated with the local and global features of the amino acids after the mutation. This combined set of features serves as the representation of the mutation. (c) Use the features of each amino acid from the partner sequence as the features for the partner itself. (d) Features of mutation and partner are sent to the backbone for feature learning by a serious of cross-attention. The MPL head processes these feature vectors through a linear layer to generate one-hot encoded classifications (disrupting, decreasing, no effect, increasing). (e) Illustrates the detailed structure of the extractor block. The features obtained from the BERT layer and injector are normalized before being sent to the cross-attention mechanism, followed by a feed-forward neural network (FFN). (f) Illustrates the detailed structure of the injector, where the output from the previous layer undergoes normalization before being processed through cross-attention.

We validated the accuracy of TAPPI through comprehensive and ablation experiments, and demonstrated the model’s interpretability and its biological relevance in areas such as reverse mutations and the distribution of coding space. Additionally, it shows promise for applications in studying long-distance multi-point mutations. We also validated that in Autism Spectrum Disorder (ASD) and two rare diseases (Lennox–Gastaut syndrome (LGS) and epileptic encephalopathies (EE)), TAPPI predicts a greater number of ‘With Effect’ trend involving partner protein that are not only closer to disease-related proteins but also more numerous compared to the control group, particularly in extreme cases (disrupting). Further enrichment analyses and case studies demonstrate the potential of TAPPI to aid in precise individual diagnosis and bridge-mutation-disease. Suggesting that the trends TAPPI predicted are associated with disease-related protein and are meaningful for studying the pathogenic processes associated with de novo mutations.

### 2.2 Performance on Quantifiable Evaluation Metrics

We will present the predictive capabilities when handling single-point mutations in this section. We calculated the predictive capabilities for single-point mutations in the test set (Fig. 2a, Table S1).

**Figure 2.**
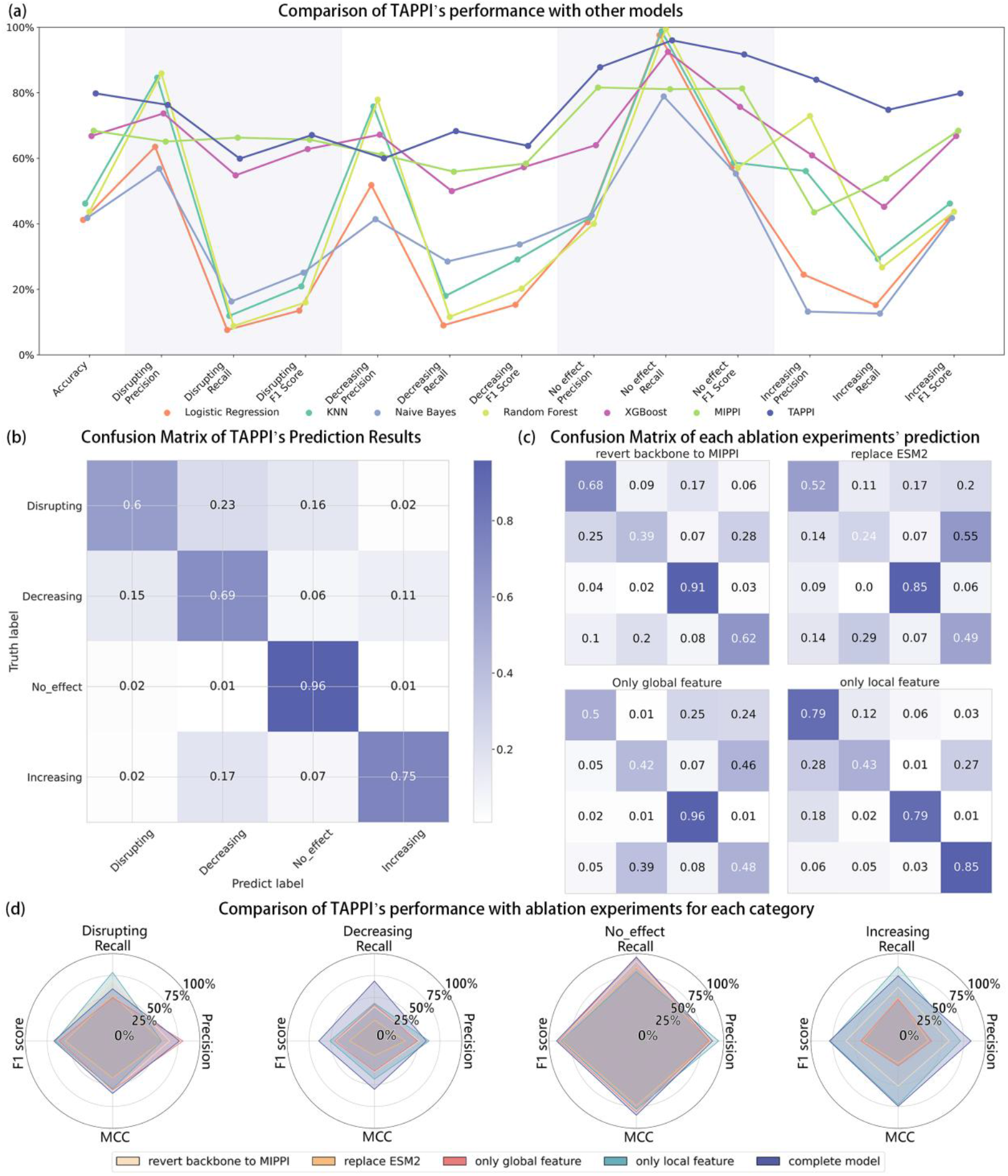
TAPPI’s performance and results from ablation experiments. (a) Comparison of TAPPI’s performance with other models in Terms of Accuracy and Metrics for Each Category: Precision, Recall, and F1 Score. (b) Confusion Matrix of TAPPI’s Prediction Results on the Test Set. (c) Confusion Matrix of each ablation experiments’ prediction results on the Test Set. (d) Comparison of TAPPI’s performance with ablation experiments for each category.

For the prediction of the impact of single mutations, our model achieved the highest scores in 9 out of 13 evaluation metrics. The accuracy for the ‘Decreasing’ class is comparable to that of MIPPI, while the recall for the ‘Disrupting’ class is lower than MIPPI. Apart from this, our model has shown notable improvements across all evaluation metrics compared to MIPPI, with the overall accuracy for the four classes increasing from 68.4% to 79.8%. Additionally, our performance in predicting the ‘Increasing’ class has also greatly improved, achieving the highest scores across all predictive performance metrics for this class. This indicates that we have resolved the issue of low prediction accuracy for the ‘Increasing’ class, enabling the model to predict ‘Increasing’ with greater accuracy. Furthermore, our model has the highest F1-score across all four classes.

Multi-point mutations are a frequent occurrence in biological organisms and are of paramount importance due to their potential to alter PPI stability. These mutations, which take place within a limited amino acid range, can lead to substantial changes in protein structure and dynamics. Such alterations are not only critical for the normal functioning of biological systems but can also be a causative factor in the development of various diseases.

The MIPPI model, while highly effective in predicting the effects of single mutations on PPI variations, has a limitation in handling multi-point mutations. This is where TAPPI comes into play, offering a sophisticated approach to tackle the complexities of multi-point mutation caused trends by containing data with multi-point mutations when training.

TAPPI is equipped with the capability to predict the collective and synergistic effects of multiple nearby mutations. By considering the interactions and potential cooperativity between these mutations, TAPPI provides a more comprehensive understanding of how such mutations can collectively influence PPI. There are 361 multi-point mutation data in the test set, with the maximum distance between mutations being within 20 amino acids. TAPPI achieved an accuracy of 67.0% for multi-point mutation prediction (Table S1), comparable to the single-point mutation prediction capability of MIPPI. Compared to the single-point mutation prediction capability of TAPPI, the predictive ability for the ‘Disrupting’, ‘Decreasing’, and ‘Increasing’ classes shows only a slight decrease; TAPPI can accurately assess these classes.

### 2.3 Improvement and Ablation Experiments

Several key improvements are used to enhance the performance of TAPPI: (1) We use ESM-2 [21] as protein Large Language model (LLM), which is trained on over 61.7 million microbial protein sequence structures providing superior prior knowledge for our analysis, to extract feature vectors from the sequence before and after the mutation, and the amino acid sequence of the partner protein. (2) Both global features and local features are important when predicting the trend [23–25]. We utilize features from a length of 20 amino acids surrounding the mutation, along with global features before and after the mutation. (3) We select a structure similar to ViT-adapter [26] as our backbone to process the feature vectors extracted from LLM, aiming for better performance in extracting features from both the mutated protein and the partner protein simultaneously to enhances the model’s ability to focus on relevant interactions, improving both the predictive performance and the interpretability of the results.

Benefiting from the improvement mentioned before, TAPPI’s prediction performance has a notable improvement and got an overall accuracy of 79.1% in the test set. We evaluated the model using the precision, recall, and F1-score for each class, as well as the overall accuracy for the four-class classification (Table S2). TAPPI performed well on the reserved 20% test set. Additionally, ablation experiments confirmed the effectiveness of these enhancements.

For the improvement of using LLM as prior knowledge, we replaced LLM with an embedding layer to validate the effectiveness of LLM. To assess the improvement of using both local and global features, we tested the model with only global features and only local features to validate the effectiveness of both. Regarding the new backbone network used in our model, we reverted the backbone to the structure that combines self-attention with convolution, as used in MIPPI, to prove its effectiveness.

We evaluate the ablation experiments using the confusion matrix of TAPPI’s predictions on the test set (Fig. 2b), along with the confusion matrices from each ablation experiment (Fig. 2c) and their corresponding quantitative evaluation metrics (Fig. 2d, Table S2). In summary, as expected, the accuracy of the ablation experiments is all below that of TAPPI. The accuracy of the four ablation experiments is as follows: 60.2%, 67.4%, 73.9%, 72.3%.

According to the ablation experiment of only using global features and only using local features for mutation proteins, the LLM features before and after the mutation site better reflect the impact of the mutation compared to the global average pooling LLM features. Additionally, using both local and global features together achieves a higher accuracy than using either one alone. According to the F1-score, using both features shows notable improvement in predicting ‘Decreasing’ and ‘Increasing’ compared to using only global features, with a substantial enhancement for ‘Decreasing’ when compared to using only local features. After replacing LLM, ‘No_Effect’ can still be accurately identified, but the performance for other categories shows a substantial decline. Similar to using only global features, replacing the backbone with MIPPI largely reduced the model’s ability to identify ‘Decreasing’ and ‘Increasing’. Almost all ablation experiment models can accurately distinguish between ‘Disrupting’ and ‘No_Effect’, but accurately identifying ‘Decreasing’ and ‘Increasing’ requires the implementation of several improvements.

Additionally, comparing the performance of MIPPI with the reverted backbone to MIPPI reveals that switching to a more advanced feature extraction method like LLM (ESM-2), MIPPI’s prediction accuracy improved over using PSSM. However, there remains a 6.9% gap in 4-class accuracy compared to TAPPI. This indicates that the backbone used in TAPPI is highly effective for predicting changes in protein binding capacity.

### 2.4 Predictive Capability of Reverse Mutation

Besides the performance of 4-class accuracy, it’s also crucial to evaluate whether TAPPI predictions correspond to real-world biological significance.

A common evaluation method is to assess the model’s ability to distinguish between forward mutations and reverse mutations caused trends, as this reflects its understanding of biological processes. To measure TAPPI’s capability in recognizing reverse mutations, we selected two evaluation approaches: first, assessing data where both forward and reverse mutations appear exclusively in the test set, and second, evaluating cases where one mutation is included in the test set while the other is present in the training set (Fig. S1).

It can be easily observed that the TAPPI’s predictive capability for reverse mutations is comparable to its overall predictive ability, showing no notable decline in performance for reverse mutations (Fig. 3e). In the four-class classification tasks for each evaluation method, each category performed well across various metrics, indicating that the model has learned biologically meaningful patterns.

**Figure 3.**
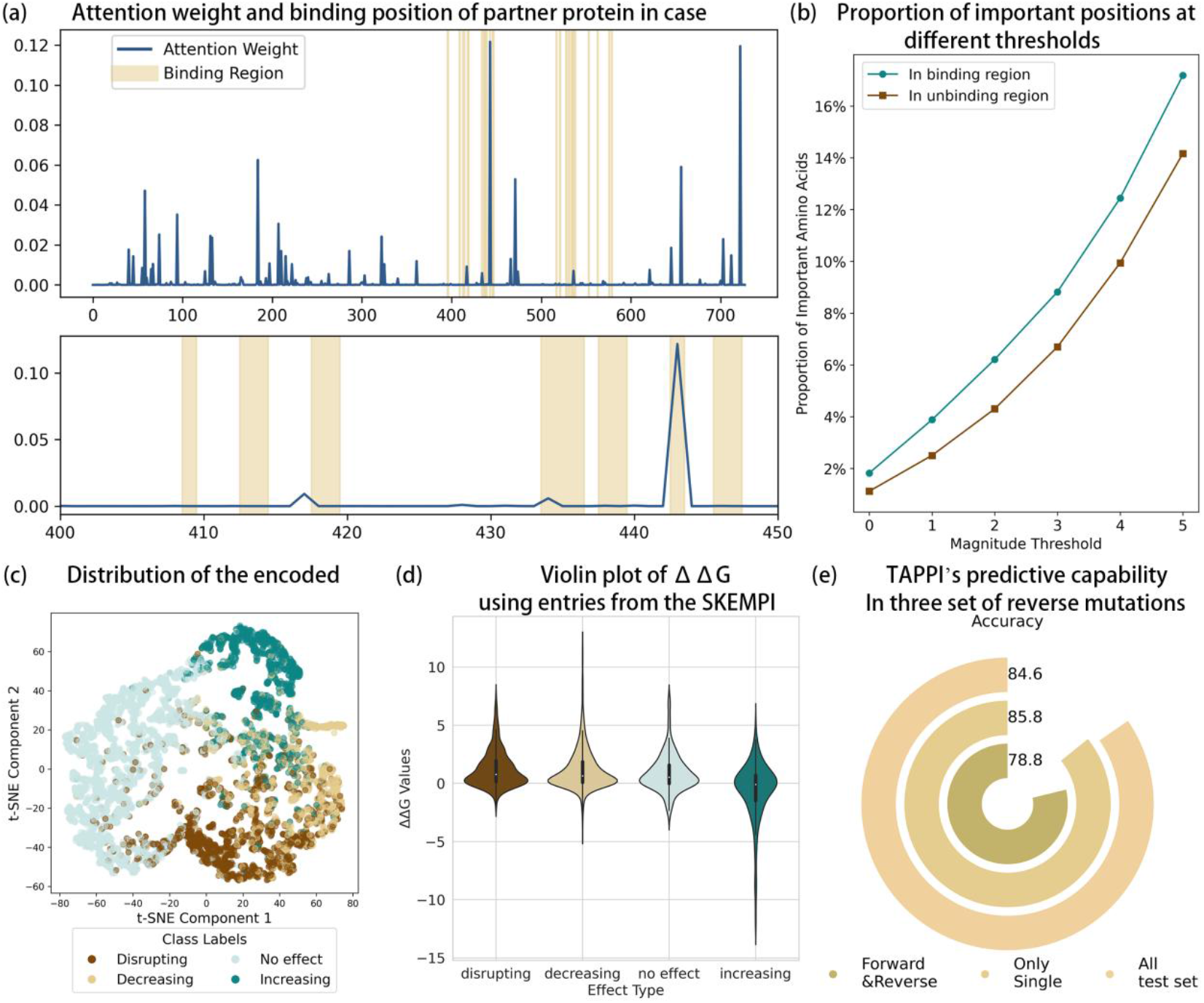
Validations of TAPPI using internal data. (a) The attention weights (dashed) for each amino acid of the partner protein, along with the original binding positions of the partner and mutations in the INSIDER database (shaded) of one ‘Disrupting’ data in the test set. (b) Proportion of important positions in binding and non-binding regions across different certification thresholds for important positions. Horizontal axis represents different evaluation thresholds for important positions; for example, a threshold of 3 indicates that positions with attention greater than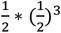 of the maximum attention value for that protein is considered important. (c) Distribution of the encoded space in the test set visualized using T-SNE. (d) Violin plot showing the distribution of ΔΔG values in different prediction categories after predicting entries from the SKEMPI v2 database using TAPPI pairs. (e) TAPPI’s predictive capability for reverse mutations in the test set is evaluated under three scenarios: (1) both forward and reverse mutations are present only in the test set (2) one mutation is present in the test set while the other is in the training set (3) all data in the test set with a label of decreasing, no effect and increasing.

However, it is important to note that the overall accuracy of the test set is fixed; thus, an excessively high accuracy in the subset could result in a correspondingly low accuracy in the complementary subset. Conversely, a low accuracy would suggest that the model struggles with reverse mutations. Therefore, achieving similar accuracy across the three datasets is an ideal outcome.

This suggests that TAPPI not only excels in predictive accuracy but also captures the underlying biological mechanisms governing protein interactions. By successfully predicting the effect of both forward and reverse mutations, the model demonstrates its relevance in biological contexts, enhancing its utility for researchers seeking to understand the implications of mutations on protein functionality.

### 2.5 Internal Validation that Reflects Biological Relevance

TAPPI is entirely based on the attention mechanism. This improvement not only improves accuracy but also provides TAPPI with excellent interpretability.

Similar to self-attention, cross-attention also possesses good interpretability due to the use of attention matrices which can be achieved by calculating the degree of attention that the results give to the features of each amino acid in (11). This allows for a clearer understanding of how different inputs influence each other within the model and an understanding of which inputs are more relevant to the global vector by analyzing the attention weights. We downloaded the ‘Highest Confidence Interfaces’ files from interactome INSIDER (http://interactomeinsider.yulab.org/) as the interface benchmark and mapped the attention weights to these PPI interfaces. These attention weights are derived from the multiplication of attention matrices across various layers, including self-attention, cross-attention, and the final attention layer that pools the feature matrix into a feature vector. These weights represent the degree of focus on each amino acid feature in the partner protein, indicating how much each amino acid contributes to the prediction outcome. This process allows for a nuanced understanding of the interactions and importance of specific amino acids in the context of the model’s predictions. We show the attention weights calculated by multiplying the attention matrix for each amino acid in a data set (Fig. 3a) to show that our model is interpretable. By statistically analyzing the number of important sites, it can be observed that PPI interfaces contain a greater number of significant positions compared to other regions (Fig. 3b).

Assessing whether the distribution of samples in the encoding space aligns with biological principles is also an important evaluation criterion for the model [27]. We use the feature vector of the test sample after pooling and before processing into one-hot encoding. T-SNE was used to reduce the dimensionality of the feature vectors to two dimensions and visualize them to observe the distribution of TAPPI’s embedding space (Fig. 3c). It can be seen that in the embedding space model can easily distinguish between ‘No_Effect’ samples and other samples. As for samples with an effect, the distribution pattern of samples ‘Decreasing’ between ‘Increasing’ and ‘Destructing’ aligns with the biological significance of gradual progression from destructing to increasing. It is also noteworthy that ‘Decreasing’ samples and ‘Disrupting’ samples are not easily distinguishable, which corresponds with real-world phenomena.

The results clearly illustrate that TAPPI can easily distinguish between ‘No_Effect’ samples and those with effects (‘Disrupting’, ‘Decreasing’, ‘Increasing’). For samples that do have an effect, the distribution pattern shows a gradation between ‘Increasing’ and ‘Destructive’ trends, which aligns with the biological significance of gradual progression. This comprehensive evaluation underscores TAPPI’s capability to capture and represent the complexities of real-world biological processes.

To further verify TAPPI’s Biological Relevance, we collected entries in the SKEMPI v2 dataset [67] with the ΔΔG labels remaining as an external validation set. We analyzed the distribution of the provided ΔΔG labels in different prediction results by TAPPI (Fig. 3d).

As the influence shifts from disrupting to increasing, the distribution of ΔΔG values shows a gradual decreasing trend. It is also noteworthy that all entries with ΔΔG less than −3.035 fall within the ‘Increasing’ category, which indicates that the mutation has a substantial positive effect on the protein’s stability and binding capabilities, potentially leading to significant changes in structure and function [28, 29]. It is also noteworthy that there is a higher proportion of large ΔΔG values in the ‘Decreasing’ and ‘Disrupting’ categories indicates that the mutation caused PPI tend not to binding anymore [28, 29].

It can also be observed that, despite the ‘Decreasing’ trend in ΔΔG values with the change in categories, which indicates that our model aligns with biological principles, this trend is not absolute. In the ‘Disrupting’ category, there are still instances where ΔΔG values are smaller than those in the ‘Increasing’ category. This suggests that our four-category classification provides a reference value that ΔΔG alone does not.

### 2.6 Key Pathogenic Proteins Related to Predicted Trend

Proteins that are closer together in the PPI network are more likely to be involved in the same KEGG pathway. If the predictions from TAPPI reflect that the PPI variation trend involving partner proteins near disease-related proteins has a higher proportion with effects, then TAPPI predicted trends that align with pathogenicity principles can assist in precise diagnosis and mechanism exploration by considering how mutations cause diseases through interfere with disease-related pathways by cause PPI variation trend.

Previous studies faced challenges when researching Epileptic Encephalopathies (EE), which is rare disease [30]. The top seven groups of proteins [31–37] related to EE, along with all human proteins and their PPIs as represented in the STRING database are collected. We statistically analyzed the proportional distribution of TAPPI predicted mutation impact categories among protein partners located at varying distances from these EE-related proteins in each individual with phenotype ‘Epileptic Encephalopathies (EE)’ and ‘Sibling Control’ from PsyMuKB database [5], and visualized the results using raincloud plots generated by the method described in Chapter 4.7 (Fig. 4 and Fig. S2).

**Figure 4.**
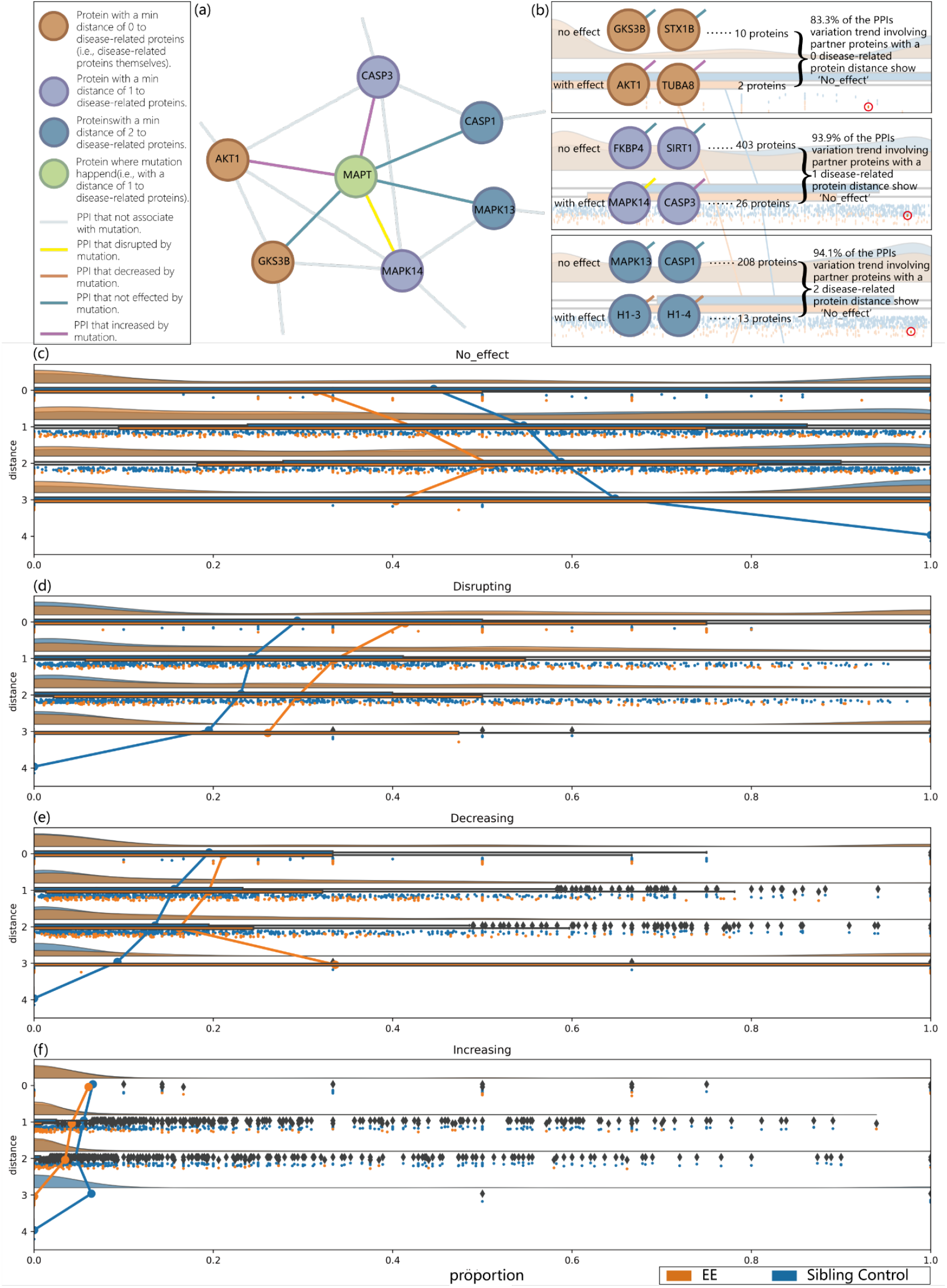
Validations of TAPPI using external data. (a) Partial PPI network of an EE patient illustrating a subset of PPI variation trend caused by the patient’s MAPT mutation (b) Illustrating all PPI variation trends of the MAPT mutation patient involving partner proteins at different disease-related-protein-distances contribute three data points with decreasing proportion as distance decreases for ‘No_Effect’ proportion statistics rain-cloud plot (c) The complete ‘No_Effect’ proportion statistics raincloud plot uses all EE and sibling control individual from the psymukb database drawn by the method described in chapter 4.7 considering the first class of proteins among the seven categories of EE-associated proteins as disease-related proteins with lessens proportion as distance lessens and more proportion in control except for extreme cases of insufficient data. (d) The complete ‘Disrupting’ proportion statistics raincloud plot with rising proportion as distance lessens and lesser proportion in control. (e) The complete ‘Decreasing’ proportion statistics raincloud plot with rising proportion as distance lessens and lesser proportion in control. (f) The complete ‘Increasing’ proportion statistics raincloud plot with rising proportion as distance lessens and more proportion in control.

In the first group of EE-related proteins, it is observed that as the distance between partner proteins and key proteins increases, there is a general trend toward higher proportions of ‘No_Effect’ (Fig. 4c), though this relationship is not strictly monotonic in extreme cases (with a small sample size at distance 3), while three categories of ‘With Effect’ decrease with a general trend toward lower proportions (Fig. 4d-f). Meanwhile, among the four trend types, raincloud plots revealed between-group differences that were more pronounced in the ‘Disrupting’ and ‘No_Effect’ categories. The patient group exhibited a higher proportion of ‘Disrupting’ and a lower proportion of ‘No_Effect’ compared to controls, highlighting the importance of predicting extreme cases (Disrupting) for disease research.

Such a phenomenon can be observed in all seven groups of EE-related proteins (Fig. S2). Moreover, the above phenomenon is not only observed in important proteins and their neighbors at distance 1, but it can also be observed at distances 2 and 3. This indicates that mutations occurring not only directly in important proteins can lead to diseases, but also indirectly lead to diseases by causing PPIs variation.

The aforementioned phenomenon indicates that (1) TAPPI can assist in precise diagnosis and mechanism exploration by concerning how mutations cause diseases through disease-related pathways, (2) the extreme cases (Disrupting), which is hard to be defined by casual PPI predicting model based on ΔΔ G, is necessary for the research on the association between mutation and disease and (3) mutations can indirectly lead to diseases by causing PPI variation trends.

### 2.7 Analysis of Individual Pathogenic Processes

As mentioned in the previous chapter, TAPPI can assist in precise diagnosis and mechanism exploration by examining how mutations cause diseases through disease-related pathways. This chapter demonstrates how to analyze the pathogenic process of mutations through KEGG pathway analysis and illustrates, through case studies, how mutations can lead to diseases either directly or indirectly.

This study performed KEGG enrichment analysis on the seven key protein groups used in the previous chapter, identifying 14 pathways with an association score > 2, calculated by average(−*log*_10_(*adjusted P* − *value*)), as associated with EE (Fig. 5a). Even the pathway, ‘Pathways of neurodegeneration’, with the lowest score among the 14 pathways can find literature supporting its association with EE [38]. Due to space limitations, we present the enrichment results of the proteins, with an trend of ‘Disrupting’, ‘Decreasing’ or ‘Increasing’ caused by mutation according to TAPPI, in each of the top 30 EE patients across these 14 pathways (Fig. 5b).

**Figure 5.**
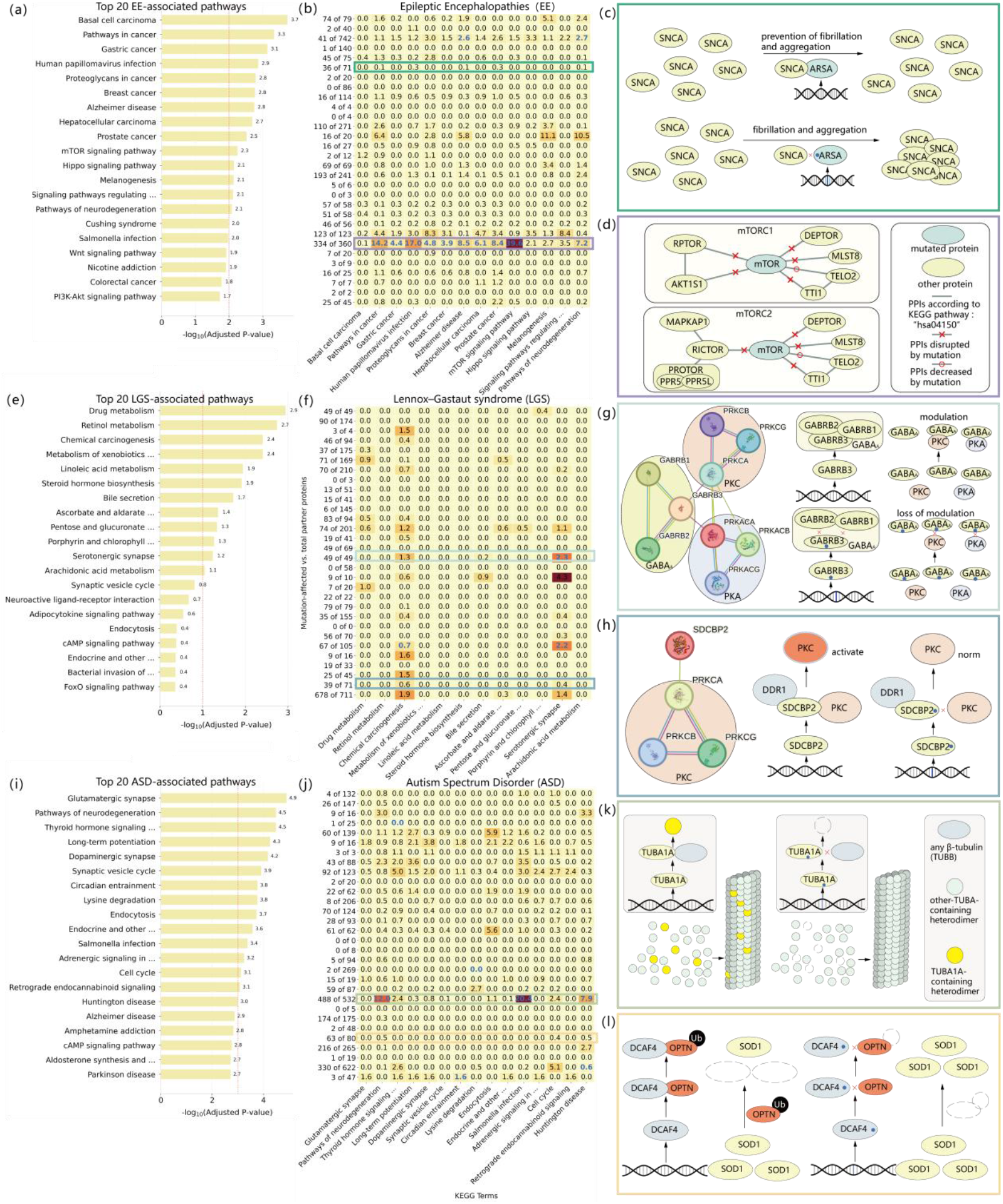
Disease-related pathway and the disturbance of them due to the PPI variation trend predicted by TAPPI. (a) KEGG pathways associated with EE and their scores calculated by average(−*log*_10_(*adjusted P* − *value*)) across seven protein group enrichment analyses. (b) Enrichment analysis of mutation-affected partner proteins from the first 30 EE patients across 14 EE-related pathways, with boxed patients further detailed in the explanation below. Blue numerals indicate mutations occurring within the corresponding pathway components. (c) Comparing wild-type ARSA preventing SNCA fibrillization and aggregation versus ARSA mutations disrupting PPI with SNCA, leading to SNCA fibrillization and aggregation. (d) Illustrating mTOR mutations impact interactions with all partner proteins in both mTORC1 and mTORC2 complexes. (e) KEGG pathways associated with LGS and their scores across six protein group enrichment analyses. (f) Enrichment analysis of mutation-affected partner proteins from the first 30 LGS patients across 12 LGS-related pathways. (g) PPI Network of GABA_A_, PKC, PKA’s components according to STRING database, with different colored edges representing the experimental methods used to discover the PPIs, and GABRB3 in GABA_A_ with PKC/PKA regulation: Wild-type function vs. mutation-induced PPI disruption. (h) The three proteins that make up PKC and their PPI with SDCBP2 and diagram comparing the wild-type process of SDCBP2-mediated DDR1 activation of PKC with how missense mutations in SDCBP2 alter its amino acid sequence, impair PKC binding by disrupting PPI between SDCBP2 and PRKCA, and consequently disrupt PKC activation. (i) KEGG pathways associated with ASD and their scores across seven protein group enrichment analyses. (j) Enrichment analysis of mutation-affected partner proteins from the first 30 ASD patients across 15 ASD-related pathways. (k) TUBA1A-TUBB Heterodimer Dynamics in Microtubule Assembly: Wild-Type Incorporation vs. Mutation-Induced Exclusion. (l) DCAF4-Mediated OPTN Ubiquitination Regulates SOD1 Degradation: Wild-Type Function vs. Mutation-Induced SOD1 Accumulation.

Among the 30 EE patients analyzed, the majority (21 patients, 70%) exhibited either direct or indirect impacts within these 14 EE-related pathways, while the remaining 9 cases require further validation using advanced methodologies. Meanwhile, only 2 of the 21 (9.52%) patients’ mutations were observed to occur directly in important pathways related to EE. This further validates the point made in the previous chapter that most mutations lead to diseases indirectly through PPIs.

Mutations in 21 patients indirectly influenced EE-related KEGG pathways through PPI. While the patient with strongest pathway perturbations carrying mTOR mutation, specifically impaired interactions with all components of both mTORC1 and mTORC2 (Fig. 5d), ultimately compromising the structural integrity of both protein complexes, which are crucial for the mTOR signaling pathway and neural signaling, and are associated with Epileptic Encephalopathies [39]. As for the least pronounced patient among the 21 patients, mutations occur in ARSA, which is not directly present in the 14 EE-related pathways, and no enrichment analysis results exceed 1 in any of these pathways. Studies demonstrate that ARSA prevents SNCA fibrillization and aggregation [40] (Fig. 5c). According to TAPPI, the least pronounced patient’s mutation cause a ‘Disrupting’ variation trend for PPI between ARSA and SNCA, resulting in uncontrolled SNCA fibrillization and accumulation, raises the risk of neurodegenerative diseases [41] and influences both ‘Pathways of neurodegeneration’ and ‘Alzheimer’s disease’ pathways. Although ARSA is not part of the two EE-related pathways, mutations in ARSA can still indirectly influence these pathways, particularly through their cause PPI variation trend with SNCA, which is a component of both pathways.

The above phenomenon further illustrates that mutations can indirectly impact key pathological pathways through causing PPI variation trend, which can be explored by TAPPI. It also demonstrates that TAPPI can assist in precise diagnosis and pathogenic mechanism exploration for the majority of patients.

### 2.8 General Applicability of TAPPI in Disease Research

To demonstrate the general applicability of our work, we also conducted the above analysis process in two other diseases.

Fully elucidating the burden that Lennox–Gastaut syndrome (LGS), a rare disease, places on individuals with the condition and their caregivers is critical to improving outcomes and quality of life [42]. Six groups of proteins related to LGS [43–48] collected from the STRING database (https://string-db.org/) were used for case analysis in LGS patients from the PsyMuKB database [5]. The results obtained are similar to those in the EE analysis, where patients exhibit more ‘Disrupting’ mutations and fewer ‘No_Effect’ mutations, indicating the feasibility of studying LGS through TAPPI (Fig. S3).

Perhaps due to the rarity of patients and the limited research available, resulting in average enrichment results, average(−*log*_10_(*adjusted P* − *value*)), of few pathways exceeding 2. Consider all pathways with a score greater than 1 as LGS-related pathways (Fig. 5e). The enrichment of the top 30 partner proteins affected in LGS patients within these pathways is shown (Fig. 5f). Twenty of the 30 patients (66.7%) were observed to have an indirect impact on these pathways through PPI.

The light-cyan-boxed patient, who scored higher in the LGS-related pathway enrichment analysis and exhibited unique direct mutations within the LGS-associated pathway, had a mutation in GABRB3, caused ‘Disrupting’ trend in PPI with GABRB1, GABRB2, and PRKCA, simultaneously affecting GABA_A_ formation and preventing PKC and PKA from binding to GABA_A_. As PKC and PKA regulate GABA_A_ function [49], impairment in the formation of GABA_A_ and its binding ability with PKA and PKC in this patient will influence the function of GABA_A_, which is responsible for inhibiting neuronal activity (Fig. 5g). This dysfunction is associated with LGS [50]. Meanwhile the deep-cyan-boxed patient, who had no enrichment analysis results greater than 1 in all LGS-related pathways, had a mutation in SDCBP2 cause a ‘Disrupting’ trend for PPI with PRKCA, present in both ‘Chemical carcinogenesis’ and ‘Serotonergic synapse’ pathways. DDR1 promotes activation of PKC via enlisting syntenin-2 (Fig. 5h) [51]. As PRKCA is the only protein among the three components of PKC that has a PPI with SDCBP2, SDCBP2 no longer binding with PRKCA will affect the activation of PKC (Fig. 5h), thereby impacting the two LGS-related pathways: Chemical carcinogenesis and Serotonergic synapse pathways. Furthermore, PKC is related to the function of GABA_A_ [49], and impaired activation of PKC will affect neuronal inhibition through its impact on GABA_A_, which is associated with LGS. TAPPI identified that SDCBP2 is not present in the ‘Chemical carcinogenesis’ and ‘Serotonergic synapse’ pathways. By causing PPI between SDCBP2 and PRKCA variation, it indirectly affects disease-related pathways.

In the analysis of Autism Spectrum Disorder (ASD) patients from the PsyMuKB database [5], similar phenomena related to EE and LGS were observed (Fig. S4) in the previously summarized seven groups of ASD-related proteins [52–58], indicates the feasibility and reliability of using TAPPI for the analysis of ASD. In the analysis of the 15 ASD-related pathways (Fig. 5i), enrichment was performed on the top 30 affected proteins of ASD patients from the PsyMuKB database (Fig. 5j). The most pronounced effect (light green box) arises from TUBA1A mutations that comprehensively cause PPI with all TUBB-class proteins, including TUBB1 and TUBB2A and seven others, disrupted. These disruption affects the formation of heterodimers containing TUBA1A, ultimately leading to a lack of TUBA1A in microtubules (Fig. 5k). Due to the relevance of TUBA1A in ASD pathophysiology [59], the absence of TUBA1A in microtubules is associated with ASD.

The orange-boxed patient, having no score greater than 1 across all 15 ASD-related pathways in enrichment analyses, occurred mutation in DCAF4, causing PPI with OPTN disrupted. As DCAF4-mediated ubiquitination of the OPTN protein promotes the autophagic degradation of SOD1 (Fig. 5l) [60], altered DCAF4 can no longer mediate the ubiquitination of the OPTN protein, which affects the promotion of SOD1 autophagic degradation, leading to the accumulation of SOD1 (Fig. 5l). The accumulation of SOD1 is associated with neurodegenerative diseases [60] and impacts the pathways of neurodegeneration, associated with ASD [61]. DCAF4, which is not part of the ASD-related pathways, indirectly affects the pathway through PPI.

These results indicate that TAPPI has broad applicability across multiple diseases, as it can assist in precise diagnosis and the exploration of pathogenic mechanisms, furthermore, highlighting the widespread indirect effects that TAPPI can uncover.

### 2.9 Discovery of Key Mutations in Long-Distance Multi-Mutations

Due to limitations imposed by the receptive field, TAPPI cannot directly handle long-distance multi-point mutations. However, it can indirectly infer the effects of multi-point mutations by sequentially predicting the impact of each individual mutation on PPI. Additionally, TAPPI is capable of identifying significant single-point mutations that have a considerable impact on PPI across various scenarios.

Additionally, 792 entries in the IMEx database contain long-distance multi-point mutations, and TAPPI can predict the individual effects of each mutation separately, allowing for the inference of the combined effects of multiple mutations, or to identify key mutations among them, thereby enhancing its utility in understanding complex protein interactions.

In the 201-long mutation protein, mutations occur at positions 42, 68, and 121, collectively leading to ‘Disrupting’ with the partner protein (Fig. S5). According to TAPPI predictions, these mutations cause ‘Disrupting’, ‘Decreasing’, and ‘No_Effect’, respectively. It is evident that position 42 is the primary factor leading to ‘Disrupting’, followed by position 68, while the impact of position 121 is relatively minor. Despite their similar attention weights on the partner protein, there are notable differences. The mutation causing ‘No_Effect’ has a lower attention weight at position 65 of the partner protein, which is the most attended position for all three mutations. Additionally, the attention weights at other positions also vary, reflecting the different roles and degrees of impact that mutations at various positions have on the partner protein in the multi-point context.

## 3. Discussion

This paper proposes TAPPI to access the protein-protein interaction (PPI) variation trend caused by mutation with state-of-the-art performance and linear complexity effectively bridging mutations and disease using discrete prediction results enabling more intuitive and clinically relevant assessments to bridge mutation and disease. In benchmark evaluations we validated the accuracy of TAPPI through comparative and ablation experiments. We also demonstrating TAPPI is interpretable and biologically relevant. In external datasets from multi disease, building upon the finding that TAPPI’s prediction is pathogenic relevant, we further demonstrate its value in enabling precise individual diagnosis and elucidating pathogenic mechanisms through case studies and the reliability of TAPPI’s bridge-mutations-disease. Additionally, it shows promise for applications in studying long-distance multi-point mutations and the pathogenic roles of mutated proteins.

Meanwhile, despite TAPPI’s accuracy and application potential, there are still many areas for improvement. Currently, TAPPI cannot directly handle the synergistic effects of long-distance multi-point interactions. Additionally, the crude concatenation of global and local features results in an underestimation of the importance of global features, weakening their ability to represent the long-distance effects of mutations. It is advisable to establish separate channels to process global features. We also believe that the classification results should include a category for partners that do not interact with the proteins undergoing mutations to enhance the model’s robustness. Furthermore, the current prediction results from TAPPI for diagnosing ‘whether a disease is present’ still require further research.

Overall, TAPPI is an accurate and interpretable framework that reveals how mutations affect disease pathways with biological relevance and potential application.

## 4 Method

### 4.1 Dataset and Pre-Processing

We use a protein mutation dataset from the IMEx Consortium [20], which features more than 50,000 annotations describing the effect of amino acid sequence changes on physical protein interactions with tested experimental evidence. We filtered out duplicate and conflicting datasets, which arose from human curation during preprocessing. The dataset is updated in the IntAct database.

Limited by the receptive field, TAPPI can only predict multi-point mutations with the maximum distance between mutations being within 20 amino acids. We removed data where the maximum distance between multiple mutations exceeded 20, resulting in 22,208 entries. Although the overhead of TAPPI does not increase with the length of the mutation protein, the overhead of LLM (ESM-2) increases quadratically with both mutation and partner lengths. To address this, we only retained data where the mutation protein length is less than 1500 and the partner protein length is less than 1000. We removed 3690 records from the 22,208 entries. The records removed represent only a small portion of the total dataset (approximately 16.6%), ensuring that the core data remains intact.

We label the three different levels of increasing as “increasing”, the three types of decreasing as ‘Decreasing’, and the three types of disruption as ‘Disrupting’, and keep the label ‘No_Effect’. We remove all ‘Causing’ data for a few ‘Causing’ class entries (only 103), and rare mutated proteins cause new PPI in nature. After the data preprocessing, we obtained a total of 18,518 entries, which include 6,657 ‘Disrupting’, 6,317 ‘No_Effect’, 4,342 ‘Decreasing’, and 1,099 ‘Increasing’.

To enhance TAPPI’s real-world relevance, we added reverse data to the dataset, a technique commonly used by other researchers [62, 63]. We generated ‘reversed entries’ by swapping labels between mutant and unmutated proteins for entries labeled as ‘Decreasing’, ‘No_Effect’, and ‘Increasing’, while considering that reversing labels for disrupting mutations lacks real-world significance. Finally, we obtained a total of 30,173 entries, which include 6,657 ‘Disrupting’, 12,634 ‘No_Effect’, 5,441 ‘Decreasing’, and 5,441 ‘Increasing’.

### 4.2 Details of ESM-2

In addition to amino acid sequences, proteins possess various physical and chemical properties that can be influenced by changes in these sequences. However, learning these complexities from a small dataset can be challenging. Therefore, incorporating prior knowledge is essential, as it can significantly enhance the performance of TAPPI. ESM-2 [21] is a large-scale protein language model trained on over 61.7 million microbial protein sequence structures. For each amino acid sequence of length *i*, after adding a start symbol and an end symbol, a feature matrix is obtained consisting of *i* + 2 vectors {*N*_0_, *N*_1_, *N*_*2*_,…,*N*_*i*+1_}. This represents that a 1280-dimensional embedding feature is obtained for the start symbol, end symbol, and each amino acid through the ESM-2 pre-trained model, each *N*_*a*_ (where *a* ∈ {1,*2*,3, . . ., *i*}) conform to *N*_*a*_ ∈ ℝ^1*2*80^, represents the features extracted from the *a*^*th*^ amino acid through the ESM-2 model within the context of the upstream and downstream amino acid sequences, including information on structure prediction, binding site prediction, mutation effect analysis, and evolutionary relationships and *N*_0_, *N*_*i*+1_represents the features of start symbol and end symbol. As the authors of ESM-2 state, averaging the vectors from the set {*N*_0_, *N*_1_, *N*_*2*_,…,*N*_*i*+1_} yields a single vector *N* ∈ℝ^1280^ that represents the global features of the protein including information on hole protein on structure prediction, binding site prediction and so on.

### 4.3 Attention Mechanism

The attention mechanism is a technique used for processing sequence data, offering strong predictive performance and interpretability. As for self-attention, when input vectors 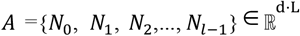 with length *L*, it computes three matrices: the query (Q), key (K), and value (V) matrices, each of length *L*. The complexity(both memory usage and time cost) for this computation is *O* (*L*·*d*), where *d* is the dimensionality of the input vectors. After obtaining these matrices, the self-attention mechanism computes the attention scores of shape *L*·*L* using the dot product of the queries and keys, resulting in a complexity of *O* (*L*^*2*^). After computing the attention scores matrix, the self-attention mechanism uses this matrix to weight the value vectors, resulting in updated features of shape *L* · *d*. In practical applications, the dimensionality *d* is often fixed, which means that the primary factor affecting computational cost is the length of the amino acid sequence *L* and the overall complexity of the self-attention mechanism can be expressed as *O* (*L*^*2*^). In practical applications, normalization, feed-forward networks (FFN), and residual connections are often added before and after the attention mechanism [64], a self-attention layer can be represented as:

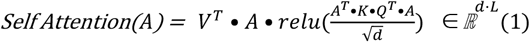

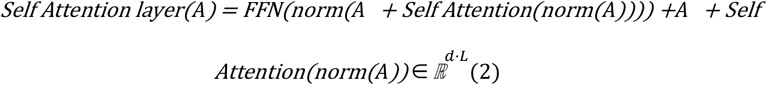

Cross-attention operates similarly to self-attention, it is used in scenarios where two different sequences are involved. Cross-attention is not only utilized in the Transformer architecture but also appears in various fields, including NLP [65] and CV [66]. When input vectors 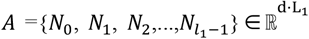 with length *L*_1_ and vectors 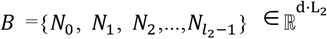 with length *L*_2_, it computes query(Q) using *A* with length *L*_1_ requires complexity *O* (*L*_1_·*d*), computes key(K) and value(V) using *B* with length *L*_*2*_ requires complexity *O* (*L*_*2*_·*d*). Then it computes the attention scores with shape of *L*_1_·*L*_*2*_. This is achieved by taking the dot product of the queries from *A* with the keys from *B*, representing the relevance of each vector in *A* to every vector in *B*, resulting in a complexity of *O* (*L*_1_·*L*_*2*_). Ultimately, the computation results in *L*_1_ vectors that represent the updated feature matrix of *A* after incorporating information from matrix *B* achieved by applying the attention weights to the values corresponding to *B*. The resulting vectors encapsulate the enhanced characteristics of *A*, reflecting the relevant features retrieved from *B* through the attention mechanism. Similar as self-attention, the complexity can be expressed as *O* (*L*_1_·*L*_*2*_). In practical application, a cross-attention layer can be represented as:

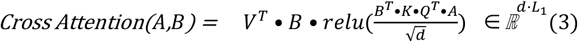

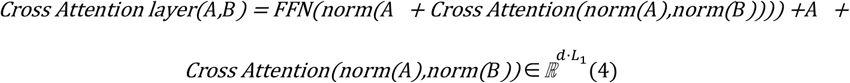

### 4.4 Backbone of TAPPI

VIT-adapter [26] is a technique in computer vision designed for dense prediction tasks. It employs multiple stacked cross-attention layers to enable interaction between the features of dense predictions and those of the Vision Transformer (ViT) [21] used in routine predictions. This integration reduces the time and memory costs associated with attention blocks when handling numerous features for dense predictions. Since ViT typically processes a smaller number of tokens derived from image patches, while dense prediction tasks involve a significantly larger number of feature vectors, the VIT-adapter efficiently manages these interactions. This disparity allows for reduced computational costs and faster processing times, making it well-suited for applications requiring extensive feature calculations in dense prediction scenarios.

The backbone of the model consists of several similar layers, and at the *i*^*th*^ layer for the input feature vectors of ViT 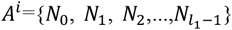 with length *L*_1_ and the feature vectors for dense prediction tasks 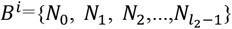 with length *L*_*2*_, the layer first use cross-attention with the query from input *A* interacts with the key and value from input *B*. This allows *A* to focus on relevant information from *B*, effectively integrating features from *B* into *A*. The result is an updated representation *A* that incorporates contextual information *B* with complexity of *O* (*L*_1_·*L*_*2*_).

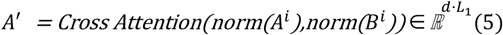

Then *A* uses its query, key, and value to perform self-attention. This process enables *A* to refine its own representation by attending to different parts of itself, enhancing its internal feature representation. It captures dependencies and relationships within *A*, improving its expressiveness for subsequent tasks with complexity of 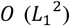.

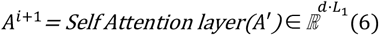

Finally the query from input *B* interacts with the key and value from the updated *A*. This allows *B* to incorporate relevant information from *A*. The result is an updated representation of *B* that reflects the context provided by *A*, ensuring that *B* is informed by the features learned from *A* with complexity of *O* (*L*_*2*_·*L*_1_).

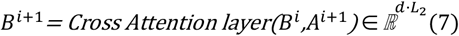

With the above process, the *i*^*th*^ layer output the input of (*i* + 1)^*th*^ layer *A*^*i*+1^ *B*^*i*+1^ enabling the stacking of multiple layers with a complexity of *O* (*L*_1_^*2*^ + *L*_1_ • *L*_*2*_).

### 4.5 Workflow

Similar to the handling of ViT vectors and dense prediction vectors, the backbone VIT-adapter can also process the vectors of mutated proteins and partner proteins by allowing the partner and mutation proteins to reference each other’s information. For data with mutation protein with length *L*_1_ and partner protein with length *L*_*2*_, we use ESM-2 to extract their feature vectors. Through ESM-2 get vectors, 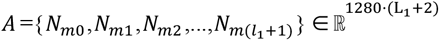 and 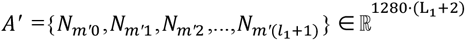 for protein before and after mutation (Fig. 1a-b), get vectors 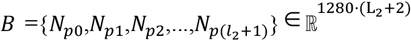 for partner protein (Fig. 1c).

For mutation protein, feature vectors of amino acid before and after mutation position(pos), an average of maximum and minimum values of the mutation position for multi-point mutation data, along with global feature vector averaging (ave) the vectors from all feature vectors are required. Although ESM-2 has already incorporated positional information for amino acids, it is necessary to add positional encoding for the mutation to distinguish between the proteins before and after the mutation. Vectors used to express mutation protein can be described as mut:

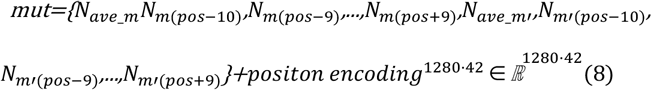

Since it’s difficult to determine the effect of the mutation on the partner (par) protein based solely on sequence information, all feature vectors of the partner protein are needed. To achieve better prediction performance and interpretability, we have added feature vectors that represent the result (res) [21] which will be updated during model training and will ultimately represent the global features of the data.

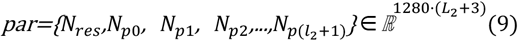

Framework of TAPPI use the global and local features of partner proteins before and after mutation calculated by (8) and features of partner protein calculate by (9) to predicts one of four possible outcomes of the mutation’s effect: strengthening the interaction (‘Increasing’), weakening the interaction (‘Decreasing’), disrupting the interaction (‘Disrupting’), or having no effect on the interaction (‘No_Effect’) (Fig 1d). To save on computational overhead, we first reduce the dimensionality of each feature vector from 1280 to 192 using a linear layer. We utilized backbone mentioned in chapter 4.4 to process these two set of features with the complexity for each layer of the model being *O* 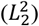 with the number of vectors for mutation within 42. Compared to the *O* (*L*^*2*^) memory usage of the self-attention layers used by MIPPI for processing partner proteins with a max length of 1000, the improved structure allows us to use more layers to enhance the model’s fitting capability. After 39 layers, we employ cross-attention to obtain the feature vector representing the outcomes in MPL head. We then use the vector to compute one-hot encoding through a linear layer for classification. Computing of the result vector can be described as:

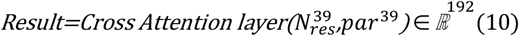

### 4.6 Interpretability of Model

An attention matrix 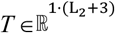 is obtained where *L*_*2*_ is the length of the partner protein, which represents the relevance between the results and the feature vectors of each amino acid in the final partner protein (Fig. 1d). In each layer of the backbone, three attention blocks can generate three attention matrices 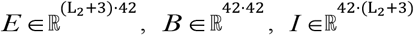 which come from the extractor block and injector. These attention matrices come from the extractor block and the injector, representing: (1) The relevance of each amino acid in the partner protein to each amino acid near the mutation sequence. (2) The relevance of each amino acid near the mutation sequence to each amino acid near the mutation sequence. (3) The relevance of each amino acid near the mutation sequence to each amino acid in the partner protein. The three matrices can be multiplied together to obtain the matrix 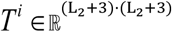. It represents the relevance of each amino acid in the partner protein to the feature vectors of each amino acid in the (*i* − 1)^*th*^ layer of the partner protein at the model’s *i*^*th*^ layer.

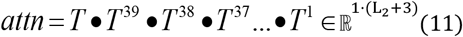

The attention score used for the interpretability studies of TAPPI is calculated by (11). This attention score represents the degree of attention that the results give to the features of each amino acid when input into the model.

### 4.7 Construction Method of Raincloud Plots

This study employed raincloud plots to visualize the proportional distribution of TAPPI-predicted partner protein categories across varying distances from disease-relevant key proteins. The protein-protein interaction network was constructed as *G* using all experimentally confirmed physical interactions from the STRING (https://string-db.org/) database (human proteome). For example, in our ASD case study, we analyzed seven distinct groups of key proteins associated with ASD. For each group of key proteins *G*_*i*_, we calculated the shortest distance from every human protein in the STRING database to that group of key proteins by (12). Here, *G*_*i*_ represents the *i*^*th*^ group of key proteins, and min_*distance* denotes the shortest path length between any two proteins in protein-protein network *G*. Furthermore, we define that if *protein*_*j*_ ∈ *G*_*i*_, *distance*(*protein*_*j*_, *G*_*i*_) = 0.

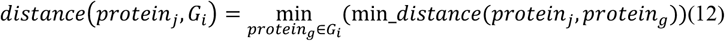

For each individual, we classified all partner proteins in group *G* of their mutated protein according to network distance as up to three partner protein group (for example, mutation happened in protein with distance 2 to *G*_*i*_, partners could have a distance of 0, 1 or 2, but partners of mutation happened in protein with distance 0 only have distance 0 or 1) and predict the proportion of effect in each partner protein group across four class of effects predicted by TAPPI. We generated 28 raincloud plots (4 classification categories × 7 key protein groups) to visualize the distance-dependent proportion of effect for each category-group combination. In the raincloud plots of the first important protein group *G*_1_ and ‘Disrupting’ in 4 classification effect, each individual contribute up to three partner protein group according to *distance*(*protein*_*j*_, *G*_1_) in (12), proportion of partner protein with ‘Disrupting’ PPI caused by mutation in each partner protein group are counted and analyzed according to different distances and phenotypes.

## Data Availability

The dataset used in training TAPPI is updated in the IntAct database (https://www.ebi.ac.uk/intact/). The “Highest Confidence Interfaces” files can be downloaded from Interactome INSIDER (http://interactomeinsider.yulab.org/). SKEMPI v2 is accessible online at https://life.bsc.es/pid/skempi2/. The protein interactions and their amino acid sequences for constructing the human PPI network can be downloaded from the STRING database (https://string-db.org/). Data for patients with EE, LGS, ASD, and sibling controls can be downloaded from PsyMuKB (https://www.psymukb.net). The KEGG pathways used can be downloaded from https://maayanlab.cloud/Enrichr/geneSetLibrary?mode=text&libraryName=KEGG_2021_Human.

## Code Availability

The code used for training, evaluating, and visualizing TAPPI can be downloaded on GitHub (https://github.com/kai3171/TAPPI).

## Acknowledgments

This work was supported by the National Natural Science Foundation of China (No: 62372099); the Jilin Scientific and Technological Development Program (No. 20230401092YY); Key Laboratory of Intelligent Rehabilitation and Barrier-free for the Disabled (Changchun University), Ministry of Education (2024KFJJ003).

